# Biofilm development of *Porphyromonas gingivalis* on titanium surfaces in response to 1, 4-dihydroxy-2-naphthoic acid - a hybrid *in vitro* – *in silico* approach

**DOI:** 10.1101/2025.06.19.660513

**Authors:** Rumjhum Mukherjee, Felix Klempt, Florian Fuchs, Katharina Doll-Nikutta, Meisam Soleimani, Peter Wriggers, Philipp Junker, Meike Stiesch, Szymon P. Szafrański

## Abstract

Colonization of titanium dental implants by the oral pathogen *Porphyromonas gingivalis* can lead to peri-implant diseases and, ultimately, implant failure. *P. gingivalis* growth can be stimulated by 1,4-dihydroxy-2-naphthoic acid (DHNA), a menaquinone precursor from various oral bacteria, yet its impact on biofilm formation remains unclear. The aim of the study was to evaluate *P. gingivalis* growth and metabolic activity over six days in response to DHNA on two titanium grade IV surfaces with different roughness using a hybrid *in vitro* – *in silico* approach. *P. gingivalis* growth was modestly stimulated by DHNA and exhibited an inverse correlation with ammonia concentration in culture medium. Notably, this growth pattern transitioned from an initial linear phase to a later exponential phase, with DHNA-treated biofilms reaching this exponential shift at an earlier stage than untreated controls. Confocal microscopy revealed that DHNA-treated biofilms exhibited surface-dependent growth patterns, with larger biofilm volumes observed on rougher surfaces in later biofilm stages, compared to smoother surfaces. Regardless of surface characteristics, the area occupied by biofilms and the size of the aggregates exhibited a consistent and progressive increase over time and was larger in late DHNA-treated biofilms. The experimental data were used to calibrate a coupled finite element method (FEM)-based model that simulated *P. gingivalis* biofilm dynamics and nutrient utilization. Summarizing, DHNA moderately stimulated *P. gingivalis* growth, accelerated its transition to ammonia-independent growth, and promoted an increase in biofilm area and aggregate size. Our coupled approach offers significant potential for advancing *in vitro* biofilm research.

**Importance:** Results of our hybrid *in vitro – in silico* experiments advance the research on *P. gingivalis* physiology and its DHNA-dependent colonization of implant surfaces. Our findings reveal that DHNA accelerates *P. gingivalis* growth, induces aggregation and promotes colonization of titanium surfaces. For the first time, DHNA-induced *P. gingivalis* growth acceleration and an earlier shift away from ammonia dependency were observed fluorometrically, highlighting ammonia assimilation as a promising marker of *P. gingivalis* physiology during early biofilm expansion. Understanding how growth factors together with surface properties influence *P. gingivalis* colonization offers a basis for future preventive strategies. Our study’s stringent characterization of 3D surface texture parameters is expected to improve reproducibility of biofilm-surface interactions experiments. The findings were validated using a continuum-based *in silico* model, initiating a hybrid approach where computational models complement *in vitro* research. Our interdisciplinary approach offers a versatile framework for investigating additional aspects of oral biofilms on titanium.

## Introduction

Titanium dental implants, valued for their durability and biocompatibility^1, 2^, face continuous microbial exposure in the human oral cavity^3, 4^ that can lead to rapid biofilm formation shortly after implantation^5^. Implant surface properties alongside bacterial characteristics drive microbial attachment, sequential colonization, and the maturation of biofilms^6–11^. These complex and inherently resistant biofilms are a major cause of recalcitrant peri-implant infections^12, 13^.

*Porphyromonas gingivalis* stands out as a prevalent, abundant, active and highly virulent species playing a pivotal role in peri-implant pathologies and serving as a valuable model oral pathogen^14–20^. This Gram-negative asaccharolytic anaerobe belongs to the Bacteroidia class and is a part of closely related trio of species collectively referred to as the red complex, which is strongly associated with clinical indicators of periodontal disease^21^. *P. gingivalis* is a highly fastidious microorganism requiring a reduced-growth environment^22^, which underscores its dependence on other biofilm members and host factors for successful implant colonization. Its initial expansion within peri-implant biofilms remains poorly understood. These processes are influenced by both beneficial interactions, such as coadhesion, enzyme complementation, food webs, environmental modifications, and cell-cell signaling, as well as antagonistic interactions, including bacteriocins, hydrogen peroxide, organic acids, low pH, nutrient competition, and bacteriophage release^23–31^. For instance, ammonium ions that are common by-products of microbial metabolism (including that of *P. gingivalis*), have been shown to play a crucial role in supporting the growth of *P. gingivalis* under certain conditions^22^.

Another such metabolite is menaquinone, or vitamin K_2_. These isoprenoid quinones are produced by a broad spectrum of species and play an essential role in electron transport, especially during anaerobic respiration^32^. Vitamin K-requiring bacteria are common, and its supplementation in growth media is a standard procedure for isolating anaerobes. Intestinal bacterial synthesis of vitamin K is also vital for animal and human nutrition. It is therefore important to understand the role of vitamin K in the ecology and physiology of the human microbiome. Menaquinones and their precursors, collectively referred to as naphthoquinones (NKs), have been identified as essential growth factors for *P. gingivalis*, exhibiting strain-dependent effects^22, 33^. Notably, approximately one-third of *P. gingivalis* genomes, including that of the type strain, are characterized by a lack of complete menaquinone biosynthesis pathway^33^. Among these compounds, the precursor 1,4-dihydroxy-2- naphthoic acid (DHNA), has been identified as potent stimulator of *P. gingivalis* growth, even in strains capable of endogenous DHNA synthesis^22^.

These findings suggest interspecies metabolic exchange as a potential driver of early *P. gingivalis* expansion^34^, a process that remains poorly understood^27^. Specifically, the release of DHNA by neighboring oral microbes may provide a critical growth advantage. Genomic profiling^3, 35^, combined with spatial mapping of oral biofilm communities^36, 37^, implicates several oral taxa as potential contributors to this interspecies metabolic support. Bacteria from several abundant oral autochthonous genera, including *Actinomyces*, *Aggregatibacter*, *Campylobacter*, *Capnocytophaga*, *Haemophilus*, *Porphyromonas*, *Prevotella*, *Rothia*, *Tannerella*, and *Veillonella*, are known producers of naphthoquinones^38–42^. Additionally, allochthonous genera such as *Cutibacterium*, *Staphylococcus*, as well as *Escherichia* and related can be abundant in certain populations and also synthesize naphthoquinones^38, 40, 43^. In humans, plasma and serum naphthoquinone levels are low, typically in the nanomolar or sub-nanomolar range, highlighting microorganisms as a key source of these compounds in peri-implant sites^44^. Notably, in the serum of pregnant women, high estrogen levels may serve as an alternative growth source, potentially reducing reliance on microbial naphthoquinones^45^. The dependency of *P. gingivalis* on metabolic interactions presents a promising avenue for antimicrobial and probiotic interventions^28, 46, 47^. However, mechanistic insights and quantitative assessments of these processes are currently lacking. For instance, the direct impact of DHNA on the growth of *P. gingivalis* on titanium surfaces has not been explored.

It is well established that bacteria, including *P. gingivalis*, exhibit enhanced colonization and biofilm formation on rougher material surfaces^48, 49^, a process that may be further facilitated by surface corrosion^50^. These findings underscore the importance of surface properties in shaping microbial physiology and influencing the outcomes in antimicrobial testing^51^. This raises an important consideration: the extent to which surface structure modulates the biofilm-promoting effects of DHNA. Given the impact of surface parameters on biofilm development, more stringent evaluation of titanium surface, such as characterization of 3D-surface texture parameters, would enhance the reproducibility of biofilm experiments in dentistry^52^. For instance, a common threshold roughness value for microbial adhesion on titanium surface (Ra < 0.2 mm) was established using contact profilometry, serving as a current standard^53^. However, advances in optical area measurement techniques have demonstrated the importance of further roughness parameters to reliably describe material surfaces, as reflected in ISO 25178. Including these variations in roughness and 3D surface texture parameters into the analysis of biofilm development would allow to further standardize and refine these approaches^54^.

One of the major challenges in oral microbial ecology research is the complexity of biofilms which hinders comprehensive experimental study. Metabolic interactions, such as ammonia or DHNA cross-feeding, which are influenced by various molecular mechanisms and shaped by physicochemical gradients, can be particularly challenging to replicate in laboratory settings. Here, advanced numerical simulations could be of major significance as predictive tools to replicate biofilm dynamics in laboratory settings^55–58^. Finite Element Method (FEM)-based modelling offers a transformative approach for biofilm research by simulating the complex, multi-scale interactions. With the ability to incorporate vast datasets and mimic environmental variables, *in silico* models can recreate the nuanced behavior of biofilms in response to various conditions, but their development require multiple iterations including validation through *in vitro* experiments.

This study examined the influence of DHNA on *P. gingivalis* biofilm formation and ammonia utilization over a six-day period on grade IV titanium disks with two distinct surface roughness profiles. Biofilm development was visualized using confocal laser scanning microscopy, and surface properties were comprehensively characterized through detailed optical three-dimensional (3D) surface texture analysis. Biofilm parameters were assessed using both univariate and multivariate statistical approaches. Our study reveals, for the first time, that small-molecule metabolic stimulation modulates *P. gingivalis* biofilm formation in a surface-dependent manner, integrating biochemical and physical surface factors. The experimental results were further integrated into a nutrient-responsive FEM-based biofilm model to enhance understanding of *P. gingivalis* biofilm dynamics and nutrient utilization.

## Materials and methods

### Culture conditions

*P. gingivalis* ATCC 22377 was cultured on Fastidious Anaerobe Agar, or FAA for short, (LAB090/NCM2020A, LabM/Neogen)) supplemented with 5% defibrinated sheep blood (SR0051E, ThermoScientific) under anaerobic conditions (80% N_2_, 10 % CO_2_ and 10 %H_2_) in an anaerobic chamber (Concept 400 Anaerobic Workstation; Ruskinn Technology Ltd., Leeds, United Kingdom) at 37 °C for 72 h.

Biofilms of *P. gingivalis* were inoculated using colony suspension method within the anaerobic chamber. Colonies from 72 hour FAA plates were washed once and re-suspended in oxygen-free Dulbecco’s Phosphate-Buffered Saline (D8537-500ml, Sigma-Aldrich) to achieve an OD_600nm_ of 0.125, corresponding to 8 · 10^7^ cells/mL and was subsequently diluted 10-fold. The purity and concentration of the inoculum were verified by plating serial dilutions on FAA. Finally, the inoculum was diluted 8-fold in culture medium to initiate biofilm formation at a starting concentration of 10^6^ cells/mL. Notably, this concentration marked the growth threshold for untreated cultures of this specific strain, while higher concentrations resulted in a diminished effect of DHNA on total growth.

### Preparation and characterization of disks

For the investigation of the 3D surface texture as a material-specific factor, grade IV titanium samples were prepared accordingly with two different roughness levels. Disks of 12 mm in diameter and 2 mm in height were finished with resin-bonded diamond grinding wheels of 45 µm grid size, later referred to as disks with “rougher” surfaces. Disks of 3 mm in diameter and 2 mm in height were polished by tumbling in suspension with 45 µm diamond polishing powder, later referred to as disks with “smoother” surfaces.

To investigate the 3D surface texture, the samples were subjected to CLSM (Keyence VK- X1000/X1050, Keyence, Osaka, Japan). A 50x objective (Nikon CF IC EPI Plan ELWD 50x; N = 0.55; WD = 8.7 mm) with a red laser (λ = 661 nm) and a resolution of 2048 x 1536 pixels was used. Twelve surfaces per sample group were measured using VK Viewer 1.1.2.174 and filtered and analyzed via MultiFileAnalyzer 2.1.3.89 (both KEYENCE, Osaka, Japan). A plane tilt for nominal form correction and an S-filter (0.8 µm; filter type: double Gaussian; end effect correction) were applied to correct the surfaces. Height (*Sa*), spatial (*Str*, *Sal*), hybrid (*Sdr*), and functional volume (*Vvv*, *Vvc*, *Vmp*, *Vmc*) 3D surface texture parameters were determined to characterize the surface as described in ISO 25178–2:2012 “Geometrical product specifications (GPS) - surface texture: areal - Part 2: Terms, definitions and surface texture parameters”. Following terminology was used:

*Sa* – arithmetical mean height (µm);
*Sdr* – developed interfacial area ratio (%);
*Str* – texture aspect ratio (Str≈0: anisotropic/directional texture; Str≈1: isotropic texture);
*Sal* – auto-correlation length (µm); *Vvv* – dale void volume (mL/m²); *Vvc* – core void volume (mL/m²);
*Vmp* – peak material volume (mL/m²);
*Vmc* – core material volume (mL/m²).

The determined parameters were further processed via IBM SPSS Statistics 29.0.0.0. The data was subjected to a Shapiro-Wilk test for normal distribution and a t-test for independent samples. The effect size was calculated using Cohen’s d. The significance level was set at 0.05.

To evaluate wettability, distilled water was applied to the sample surfaces using the DSA25S Drop Shape Analyzer (KRÜSS GmbH, Hamburg, Germany). The volume was progressively increased in 0.2 µL intervals according to the advancing-contact-angle analysis and the contact angle was measured at 23°C after 20 seconds waiting time on both sides using ellipse as fitting method^59^.

### Biofilm development assays on implant materials

All titanium disks were autoclaved prior to use. 3 mm and 12 mm disks (with smoother and rougher surfaces, respectively) were placed in each well of 96-well microplates (#267578, Nunc^TM^ Edge^TM^ 2.0 96-well, Non-Treated, Flat-Bottom Microplate) and 24-well plates (#662160, Greiner Bio-one, CellStar, MultiWell plates, PS, 24 well, lid with condensation rings, sterile, 100/PAK), respectively. Cultures of *P. gingivalis* strain was inoculated into Brain Heart Infusion broth (BHI in short, CM1135, Oxoid) supplemented with 0.4 mg/mL cysteine and 5 mg/L hemin (BHI_CH, in short). For the experimental groups, 1,4-dihydroxy-2-naphthoic acid (DHNA) was added to achieve a final concentration of 6 µM. The multi-well plates were incubated for 1 – 6 days in similar anaerobic conditions as the primary culture, at 37 °C. Negative controls comprised of disks in medium without bacterial inoculum, while positive controls contained *P*. *gingivalis* cells in BHI_CH medium, without DHNA supplementation.

### Measurement of total bacterial growth

Total bacterial growth was determined as the combined growth of the non-attached planktonic phase and the biofilm phase in multi-well plates without titanium disks. To assess the total growth of *P. gingivalis* in DHNA-supplemented BHI_CH medium, we measured optical density at 600 nm (OD_600nm_) using a Tecan multiplate reader (Tecan Infinite® M200 Pro Multi Mode Microplate Reader Fluorescence Luminescence).

### Estimation of ammonia concentration

Nutrient utilization by biofilm-forming bacteria was assessed by measuring ammonium ion concentrations in the bacterial growth supernatant using the Merck Ammonia Assay Kit (MAK310- 1KT). The assay relies on the o-phthalaldehyde (OPA) reaction, where OPA reacts with ammonium ions to produce a fluorescent compound (λ_ex_ = 360 nm, λ_em_ = 450 nm), with fluorescence intensity directly correlating to ammonium concentration. Briefly, 10 µL of standards and samples were added to 96-well plates, followed by 90 µL of a Working Reagent Mix (Ammonia Assay Buffer, Reagent A, and Reagent B), then incubated in the dark for 15 minutes at room temperature. Fluorescence was measured (λ_ex_ = 360 nm, λ_em_ = 450 nm) using a Tecan multiplate reader and ammonium concentrations were determined by comparing sample intensities to a standard curve generated with NH_4_Cl standards provided in the kit. Two technical replicas were used. The kit has a linear detection range of 0.012 – 1 mM ammonia in a 96-well format.

### Biofilm examination with confocal laser scanning microscopy

After gentle removal of the supernatant from the wells and careful transferring of the disks into 6-well plates (#657160, ThermoFischer Greiner Bio-one Polystyrene 6-well Multiplates), the biofilms were stained with the LIVE/DEAD BacLight Bacterial Viability Kit (Invitrogen), rinsed, and then fixed in a 2.5% glutaraldehyde solution prepared in phosphate-buffered saline. Confocal laser scanning microscopy (CLSM; Leica TCS SP8, Leica Microsystems, Mannheim, Germany) was employed to examine the biofilms using 40x objective lens, following the methods outlined in Dieckow et al.^60^. A 488 nm laser with an emission range of 500–545 nm was used to detect SYTO9, while a 552 nm laser with an emission range of 590–680 nm was used for propidium iodide (PI). A total of 160 biofilm images (83 on smoother surfaces and 77 on rougher surfaces) were captured, out of which 80 images were of biofilms grown without DHNA supplementation in the medium and the remaining were of biofilms grown with DHNA supplementation in the medium. At least three representative image stacks, each with optical sections of 3 µm, were acquired from each examination surface. Image analysis and 3D reconstructions were conducted using Imaris Cell Imaging software (Imaris x64, 6.2.1, Bitplane AG, Zürich, Switzerland), with biofilm volumes calculated through the surface wizard setting. Green (SYTO9), red (PI), and yellow (co-localized SYTO9 and PI) fractions were quantified. Volumes were classified as containing non-permeable viable cells when green signals were present without accompanying red signals. Parameters including the number and mean size of microcolonies, total area covered, and percentage of area covered were computed using ImageJ software v1.48 (Wayne Rasband, National Institutes of Health, Maryland, USA). Specific parameters, *i.e*., biofilm viability, biofilm volume, biofilm-covered area, micro colony/aggregate count, and mean micro colony/aggregate size were included in subsequent analyses.

### Multivariate Analysis of Biofilm Profiles

Analyses were conducted using PRIMER version 7 with PERMANOVA+ (an add-on package that extends PRIMER’s methods)^61, 62^. Five biofilm parameters were analyzed: biofilm viability, volume and area covered, as well as aggregate count and mean size. Scatter plots of variable pairs were created to explore the relationships between biofilm parameters, followed by correlation analysis using the ‘Draftsman Plot’ routine. To ensure equal weighting, each biofilm parameter was scaled so that its maximum value across all biofilm profiles was set to 100, using the ’Standardize by Maximum’ routine. Non-metric multidimensional scaling (nMDS) was performed using the Euclidean distance matrix of biofilm profiles, either for all individual samples or, for better clarity, averaged by experimental group and time point. Vector overlays were applied to examine correlations between biofilm parameters and the ordination axes. Each vector originated at the center (0,0) and extended to coordinates (x,y), representing the Pearson correlation coefficient between the variable and the first two ordination axes. The vector’s length indicated the strength, while its direction showed the nature of the relationship between the variable and the axes. Permutational MANOVA (PERMANOVA) was used to evaluate the simultaneous response of biofilm profile Euclidean distances to three factors: DHNA supplementation, surface type, and time. Type III sums of squares (partial), with fixed effects summing to zero for mixed terms, were used, along with unrestricted permutation of raw data and 9999 permutations. PERMANOVA was followed by Distance-based test for homogeneity of multivariate dispersions (PERMDISP).

### Simulations; kinematics

The deformation gradient maps a point **X** from the reference configuration Ω_0_ to the current configuration Ω_t_. This mapping is split into a growth and an elastic part, such that **F** = **F**_e_ **• F**_g_, where **F**_g_ maps a point **X** onto an intermediate, stress-free configuration Ω_g_. The growth part is modelled with the local expansion parameter α in the form **F**_g_ = α **I**, with the identity tensor **I**. The local expansion parameter evolves based on the evolution equation stated below. The intermediate configuration may not be compatible, i.e. overlapping or holes might occur. This compatibility issue is resolved by the elastic part of the deformation gradient **F**_e_, which then maps from the intermediate configuration onto the current configuration. The current configuration Ω_t_ is the physically sound, grown state of the biofilm. From the elastic part of the deformation gradient **F**_e_ the elastic right Cauchy-Green deformation tensor and its isochoric form derive:

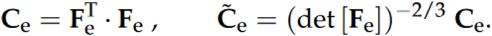

### Simulations; coupled equations and material model

As presented in our previous work^58^, the extended Hamilton principle for biofilm growth with the assumption of a quasi-static and isothermal process is formulated as

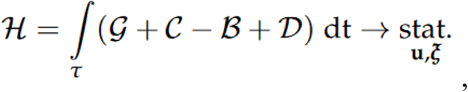

where 𝝃 and **u** represent the vectors of internal state variables and displacement components respectively and with the total potential

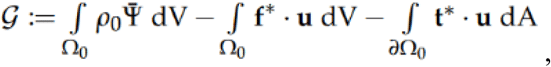

the constraint functional

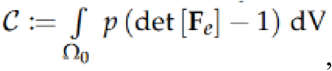

the biological work

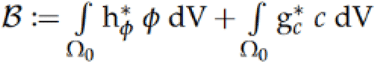

and the dissipative energy

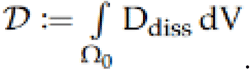

The dissipative energy D_diss_ is modeled as 𝑫_𝒅𝒊𝒔𝒔_ ≔ 𝐩^𝒅𝒊𝒔𝒔^ ⋅ 𝝃. The non-conservative forces are computed as 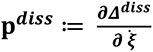 with the dissipation function, which is modeled as rate dependent, 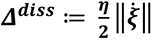. Further information on the modeling of the dissipative function can be found in our previous work^63^. To model the material behavior, an appropriate energy density function has to be chosen. The energy density function^58^

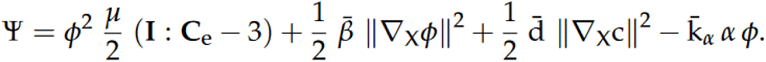

shows a good qualitative agreement with the growth of biofilm. The growth behavior, namely the terms h*_φ_ and g*_c_ inside the term describing the biological work, is modelled as

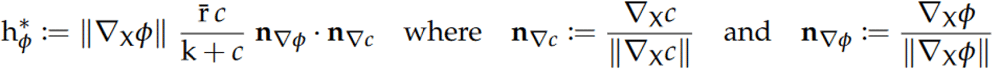

and

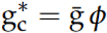

Evaluation of the stationarity conditions of H leads, along to the balance of linear momentum in its weak form, to the evolution equation for the biofilm indicator variable φ, the nutrients variable c and the variable describing the growth part of the deformation gradient α as shown below

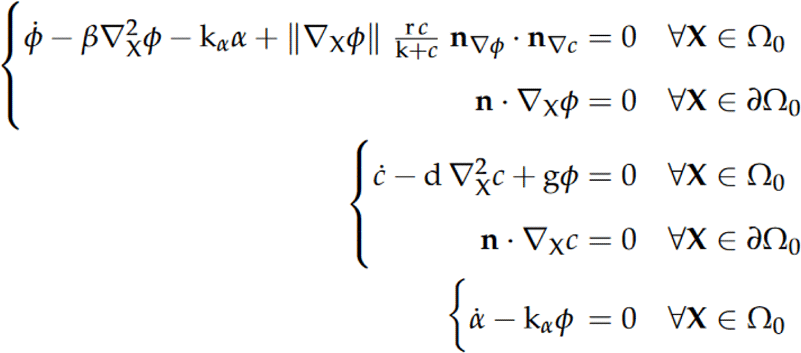

For a more comprehensive derivation of the material model, the interested reader is referred to our previous work^58^.

### Simulations; a numerical implementation

The mathematical model was solved with a standard Galerkin Finite Element Method^64^. An eight-node brick element with five unknowns per node was utilized to solve the coupled equations. The unknowns are the displacements in all three spatial directions **u** = (u,v,w)^T^, the concentrations of nutrients c and the biofilm state variable φ. The hydrostatic stress p is approximated with C^0^ ansatz functions, *i.e.*, it is approximated with spatially constant values in each finite element.

## Results

### DHNA accelerated total growth of P. gingivalis and its ammonia utilization at an early stage

To evaluate the general growth, cultures of the type strain *P. gingivalis* ATCC 22377 were measured over time. We hypothesized that assessing biofilm structure, beyond biomass, is key to capturing the effects of DHNA. As shown in the **Fig. S1**, DHNA’s impact is most pronounced at lower inoculum sizes, with 10^6^ cells/mL selected as the optimal level to study biofilm development near the stimulation threshold. Subsequently, we assessed the effect of medium richness on total growth, with and without various DHNA concentrations, across four passages over nearly 50 days (**Fig. S2**). A stimulatory effect of DHNA at concentrations between 0.12 and 1.25 mg/L was observed in full- to quarter-strength media over three passages, and persisted longer at the highest DHNA concentrations. Full-strength medium and 1.25 mg/L of DHNA were used in all following assays.

In subsequent experiments we monitored both growth and the levels of ammonia in culture conditioned medium (**Fig. 1**). Total growth increased consistently over time (**Fig. 1a**). DHNA slightly induced the total growth of *P. gingivalis* in early stages with a significant effect observed on day 3. In presence of DHNA, biofilms depleted ammonia more with a significant effect observed on day 2 (**Fig. 1b**). When correlating these results, we observed a linear relationship between growth and ammonia utilization in an early biofilm phase (2 – 3 days) that shifted after depletion by approximately 4 mM ammonium concentration to an exponential pattern in later stages (**Fig. 1c** and **1d**), which suggests the dependency on ammonium for early *P. gingivalis* growth. Interestingly, DHNA shortened this early phase by one day (**Fig 1c** and **1d**).

**Figure 1.**
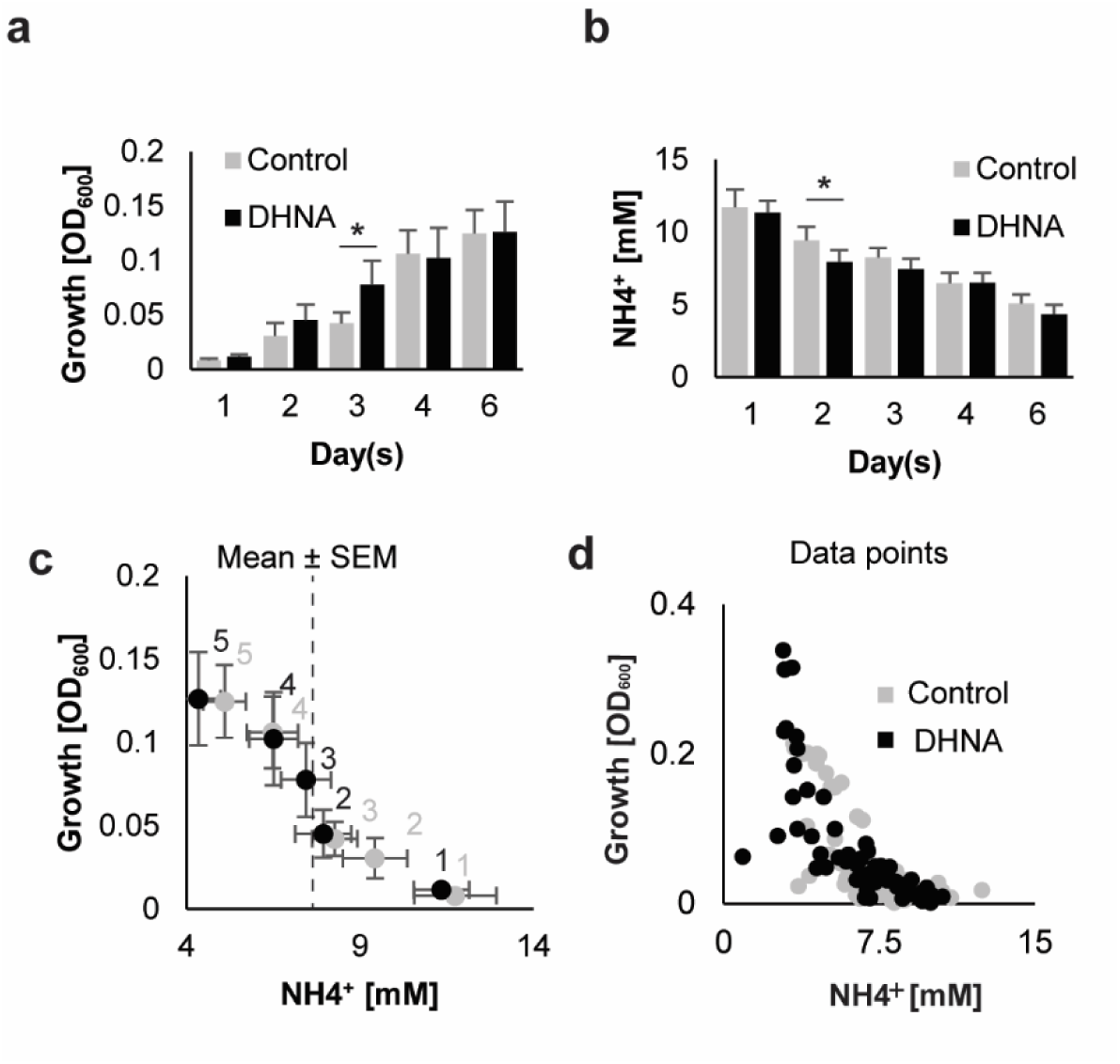
Culture growth and ammonia depletion. **a**, Total growth of *P. gingivalis* over 6 days without (control; n = 4 biological, ≥ 12 technical) and with 6 µM DHNA (treatment; n = 4 biological, ≥ 12 technical). **b**, Ammonia concentration in *P. gingivalis* cultures over time. **c**, Relationship between ammonia concentration and *P. gingivalis* growth. Mean ± sem showed. Single-tailed paired t-test with Benjamini-Hochberg correction. * for adjusted *P* < 0.05. **d.** Individual data points for total growth of *P. gingivalis* cultures over 6 days and the corresponding ammonia concentration in absence (control) and presence of 6 µM DHNA.

### The distinct surface textures aid in studying effects of surface roughness on *P. gingivalis* **biofilms.**

In our study, we explored how DHNA influences biofilm development of *P. gingivalis* over six days, using grade IV titanium disks with varying surface roughness. The foundation of this research lies in a rigorous characterization of the two surfaces, categorized as “rougher” and “smoother” (**Figs. 2a** – **2c**, **Tab. 1**).

**Figure 2.**
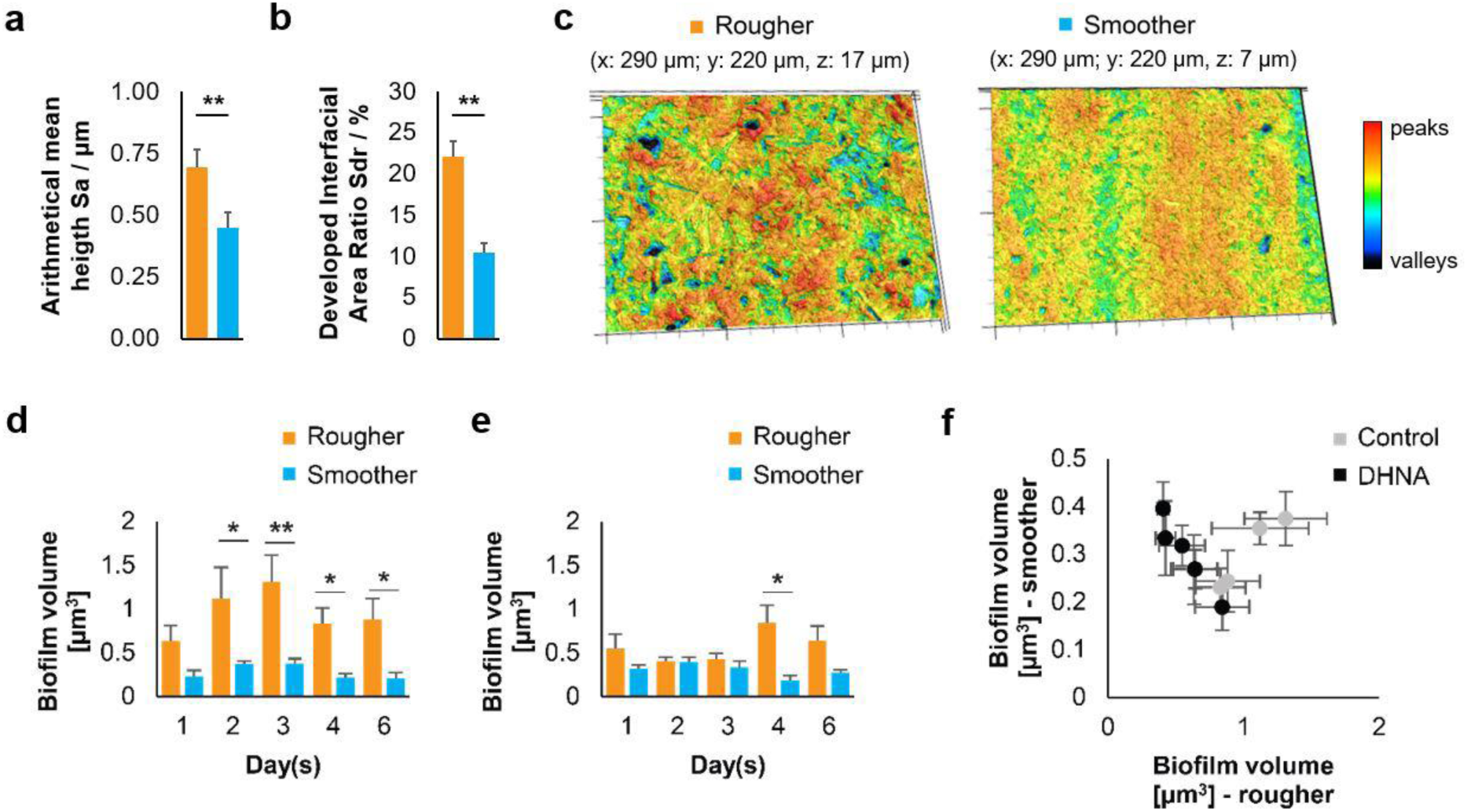
Volumes of biofilms on two titanium surfaces with different roughness (rougher and smoother). **a**, arithmetical mean heigth for two surfaces. **b**, developed interfacial area ratio for two surfaces. **c**, representative confocal micrographs of two surfaces. **d**, Volumes of biofilms on two titanium surfaces without DHNA (control; n = 4 biological, ≥ 12 technical). **e**, volumes of biofilms on two titanium surfaces with 6 µM DHNA (treatment; n = 4 biological, ≥ 12 technical). **f**, relationship between biofilm volume on two surfaces: rougher and smoother. Mean ± sem showed. Two-tailed unpaired (a and b) and single-tailed paired (d and e) t-test with Benjamini-Hochberg correction. * for adjusted *P* < 0.05, ** for adjusted *P* < 0.01.

**Table 1.**
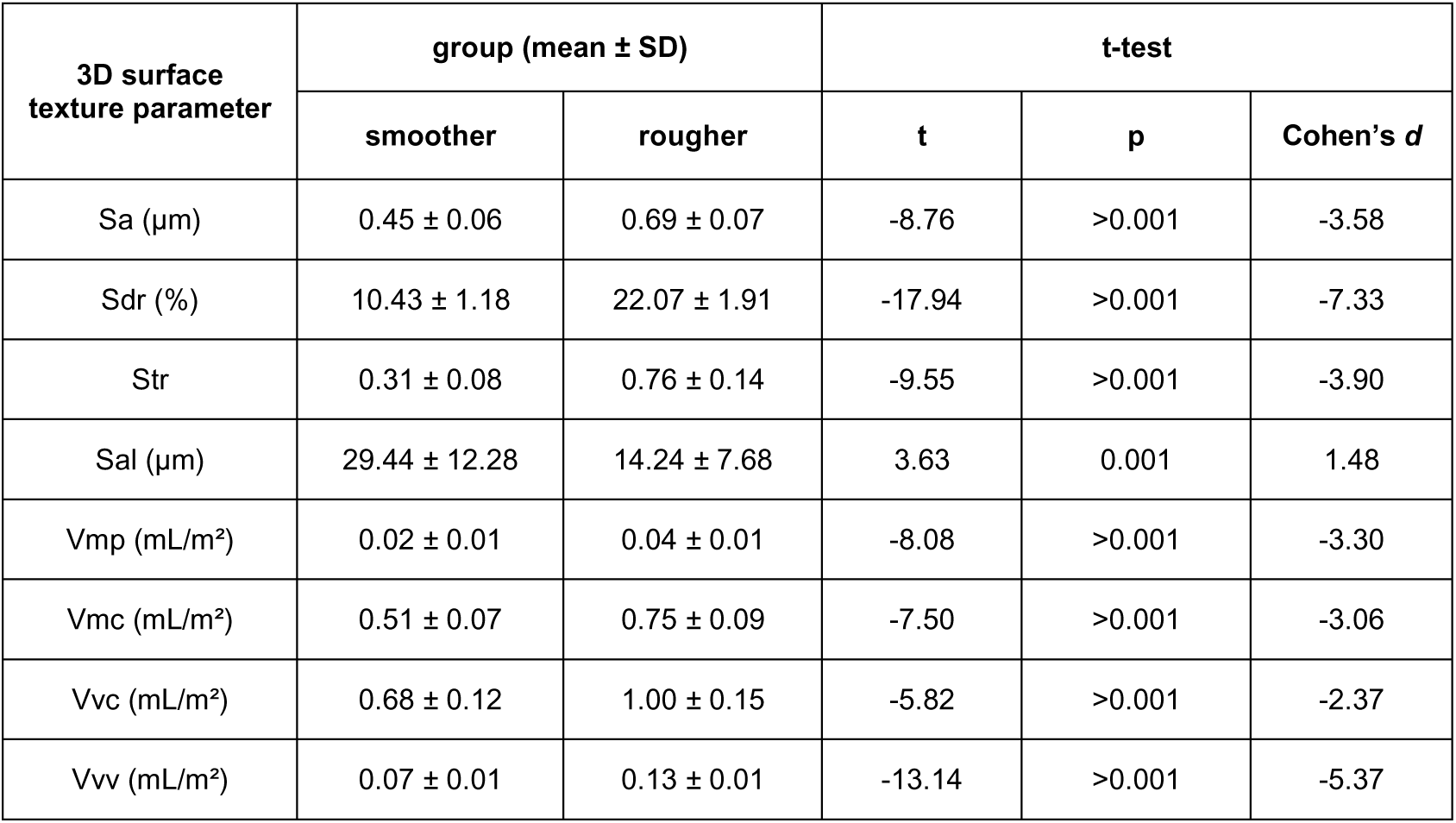
3D surface texture parameters for the investigated surfaces.

| 3D surface texture parameter | group (mean ± SD) | | t-test | | |
| --- | --- | --- | --- | --- | --- |
|  | smoother | rougher | t | p | Cohen's *d* |
| Sa ($\mu\text{m}$) | 0.45 ± 0.06 | 0.69 ± 0.07 | -8.76 | >0.001 | -3.58 |
| Sdr (%) | 10.43 ± 1.18 | 22.07 ± 1.91 | -17.94 | >0.001 | -7.33 |
| Str | 0.31 ± 0.08 | 0.76 ± 0.14 | -9.55 | >0.001 | -3.90 |
| Sal ($\mu\text{m}$) | 29.44 ± 12.28 | 14.24 ± 7.68 | 3.63 | 0.001 | 1.48 |
| Vmp (mL/m<sup>2</sup>) | 0.02 ± 0.01 | 0.04 ± 0.01 | -8.08 | >0.001 | -3.30 |
| Vmc (mL/m<sup>2</sup>) | 0.51 ± 0.07 | 0.75 ± 0.09 | -7.50 | >0.001 | -3.06 |
| Vvc (mL/m<sup>2</sup>) | 0.68 ± 0.12 | 1.00 ± 0.15 | -5.82 | >0.001 | -2.37 |
| Vvv (mL/m<sup>2</sup>) | 0.07 ± 0.01 | 0.13 ± 0.01 | -13.14 | >0.001 | -5.37 |

Our analysis revealed significant differences between surfaces in all investigated 3D surface texture parameters (**Tab. 1**). The rougher surface was initially characterized with a Sa of 0.69 ± 0.07 µm (**Fig. 2a**), showed no direction-dependent texture (Str: 0.76 ± 0.14; Sal = 14.24 ± 7.68 µm) and indicated a high specific surface area (Sdr = 22.07 ± 1.91%, **Fig. 2b**). The smoother surface showed a Sa of only 0.45 ± 0.06 µm (**Fig. 2a**), a direction-dependent texture (Str: 0.31 ± 0.08; Sal: 29.44 ± 12.28 µm) and a lower increase of the specific surface area (Sdr = 10.43 ± 1.18%, **Fig. 2b**). The functional volume parameters also showed differences between the two surfaces. Rougher surface exhibited a higher material fraction for peaks (Vmp: 0.04 ± 0.01 mL/m²) und core surface (Vmc: 0.75 ± 0.09 mL/m²) in comparison to the samples with smoother surface (Vmp: 0.02 ± 0.01 mL/m², Vmc: 0.51 ± 0.07 mL/m²). The available volumes for wetting the surface were also higher for the rougher surface (Vvc: 1.00 ± 0.15 mL/m², Vvv: 0.13 ± 0.01 mL/m²), than for the smoother surface (Vvc: 0.68 ± 0.12 mL/m², Vvv: 0.07 ± 0.01 mL/m²). The comparison of the parameters using a t-test revealed different effect sizes, according to which the effect for the comparison of Sdr (Cohen’s d: -7.33, **Fig. 2b**) and Vvv (Cohen’s d: -5.37) were the largest (**Tab. 1**). The wetting of the titanium disks with water yielded no quantifiable results. The surfaces were wetted to such an extent under 20 seconds that a valid analysis of the contact angle was no longer possible. In summary, the two surfaces exhibited significant differences in their 3D texture, making them a valuable tool for investigating the impact of surface properties on *P. gingivalis* biofilm development (**Fig. 2c**).

### Volumes of P. gingivalis biofilms varied based on surface roughness

*P. gingivalis* biofilm development was analyzed on these two surfaces by CLSM. Within the control group without DHNA supplementation, biofilm volumes reached their maximum on day 3 on both surfaces and were generally 2- to 3-fold higher on the rougher surface, compared to the smoother surface throughout the experiment (**Fig. 2d**). This difference may be attributed to the rougher surface providing more area and volume for colonization. In DHNA stimulated biofilms, volumes were significantly higher on the rougher surface only after longer incubation time (**Fig. 2e**). Biofilm volumes in the control group showed similar dynamics across both surfaces, while those in the DHNA-treated group surprisingly exhibited an inverse relationship (**Fig. 2f**).

### Surface covered by biofilm and aggregate size of *P. gingivalis* both increased in presence of DHNA

*P. gingivalis* biofilms remained consistently viable on both smoother and rougher titanium surfaces over the 6-day experiment, irrespective of DHNA treatment (**Fig. 3a**). Dynamics of volumes was similar for control and DHNA-treated biofilms on smoother surface but different on rougher surface (**Fig. 3b**). On the smoother surface, the biofilm volume increased, irrespective of DHNA treatment, in the first 2 days, before reducing in the later developmental stages. In contrary, on the rougher surface, the volume of untreated control biofilms peaked on the third day, followed by a decrease in the later stages. For DHNA-treated biofilms on the same surface, biofilm volumes slightly decreased from the first day to the third day, then increased to a peak on the fourth day, followed by a reduction on the sixth day. So, a consistent trend was not observed in case of DHNA-treated biofilms on the rougher surface.

**Figure 3.**
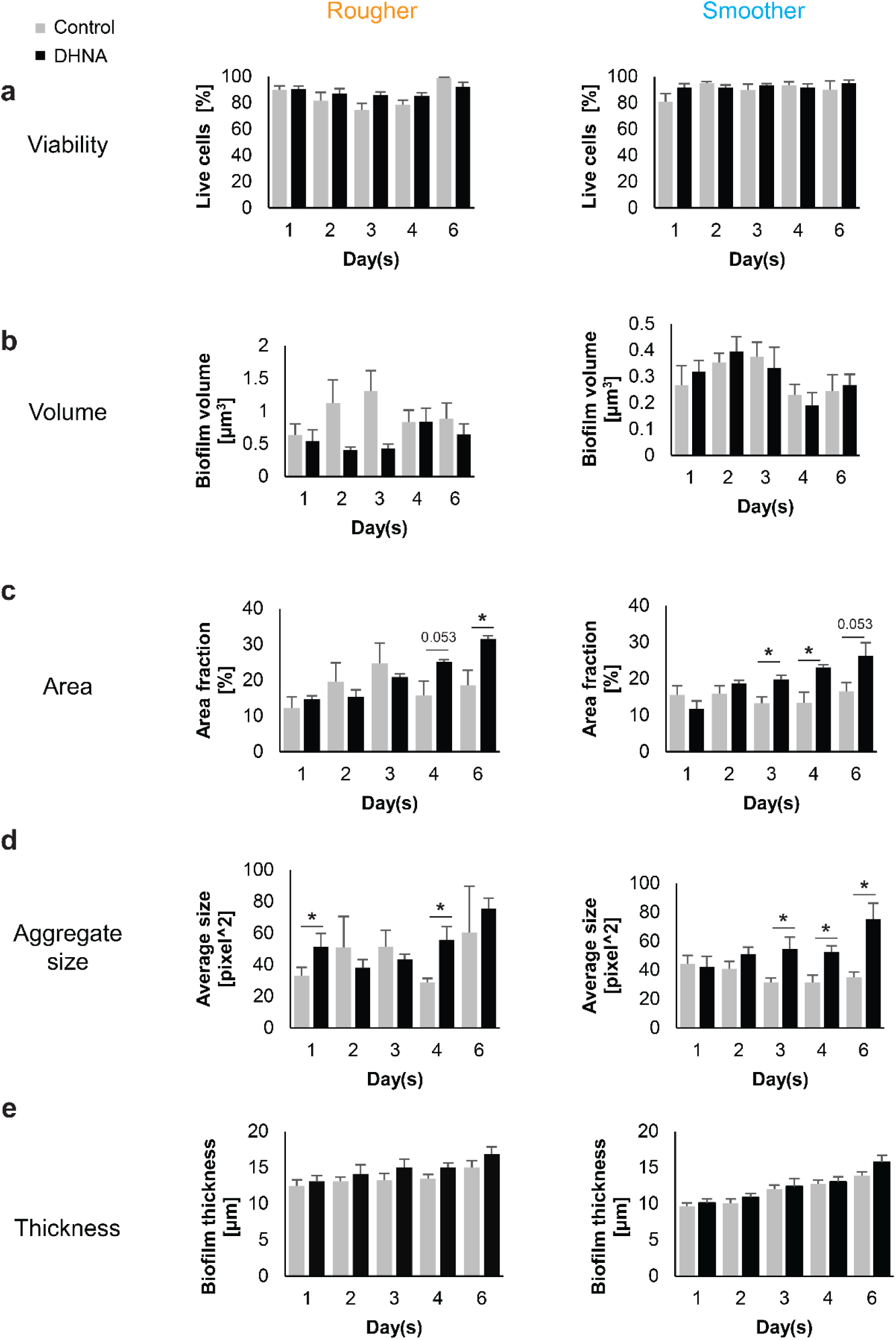
Dynamics of biofilm parameters in response to 6 µM DHNA supplementation and surface type. **a**, viability. **b**, volume. **c**, surface area covered. **d**, aggregate size. **e.** biofilm thickness. Four experimental groups represent combinations of DHNA absence/presence and rougher/smoother surface (n = 4 biological, ≥ 12 technical each). Mean ± sem showed. Single-tailed paired t-test with Benjamini-Hochberg correction. * for adjusted *P* < 0.05. Borderline *P* values are indicated on the plots.

Regardless of surface properties, the surface area covered by DHNA-treated biofilms expanded consistently over time (**Fig. 3c**). On rougher surfaces, DHNA-stimulated biofilms covered on average 31.5% of the area by day 6, roughly 1.5 times the coverage of control biofilms (18.5%). On smoother surfaces, DHNA-treated biofilms increased coverage from middle to late biofilm stages, reaching 26% compared to 16.6% in controls by day six.

Aggregate size in DHNA-stimulated biofilms increased steadily on both surfaces, peaking higher than control biofilms (**Fig. 3d**). On rougher surfaces, DHNA-treated biofilms formed larger aggregates during early and mature biofilm phases (24 h and later developmental stages), while on smoother surfaces, the aggregate size increased consistently during biofilm maturation.

Regardless of DHNA treatment, biofilm thickness increased progressively over time on both surfaces (**Fig. 3e**). Compared to the control, the DHNA treatment consistently resulted in a slightly greater biofilm thickness at each time point.

### Effect of DHNA treatment, surface roughness and cultivation time on *P. gingivalis* biofilms captured with multivariate analysis

We conducted a multivariate analysis of biofilm profiles using five aforementioned parameters (biofilm viability, volume and area covered, as well as count and mean size of aggregates). The highest correlation of 0.67 among the parameters was observed between biofilm area coverage and mean aggregate size. This trend remained consistent when smoother and rougher surfaces were considered separately, with correlations of 0.62 and 0.69, respectively. Analysis by PERMANOVA showed that surface had the greatest impact on biofilm profiles, followed by time and DHNA treatment (**Tab. 2**). The effect of DHNA was time-dependent and high residual variation was observed. When surfaces were analyzed separately, the time effect and high residual variation persisted, with the time effect being notably stronger on the rougher surface (**Tab. S1**). Interestingly, DHNA treatment had a markedly stronger effect on the smoother surface compared to the rougher one, and only on the smoother surface was this effect time-dependent. On the rougher surface, biofilm profiles were highly dispersed (F = 16.6, p = 0.0004), whereas no such dispersion was observed on the smoother surface (F = 0.1, n.s.). Ordination of treatment-surface combinations showed both spatial and dispersion effects (**Fig. 4a**). A second ordination, summarizing treatment-surface-time combinations, provided a more comprehensible overview, with point positions indicating unique dynamics for each group (**Fig. 4b**). DHNA-induced shifts were observed on both surfaces, with biofilms on the rougher surface showing greater dispersion.

**Figure 4.**
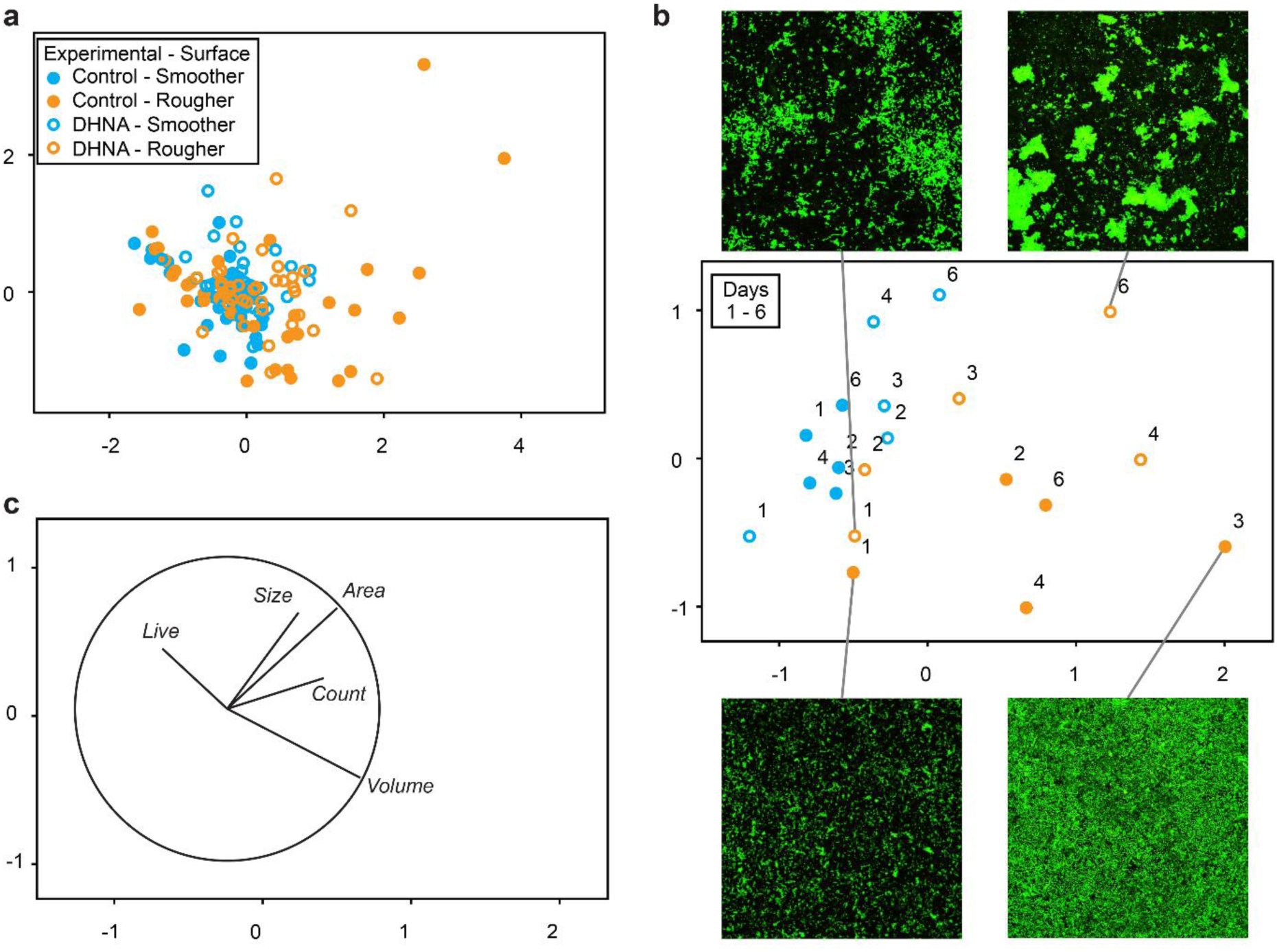
Changes in biofilm profiles in response to 6 µM DHNA supplementation and surface type. Multivariate analyses were performed. **a**, ordination of all 200 generated biofilm profiles spanning control and treatment groups (without and with DHNA), 2 surfaces, and 5 time points. Data represent 4 biological and ≥ 10 technical replicas. Five biofilm parameters, *i.e.*, biofilm viability (live), biofilm volume, area covered, aggregate size and aggregate count were standardized by maximum prior to pairwise calculations of Euclidean distances between biofilm profiles. 2D stress was 0.12. Symbols and colors indicate combinations of DHNA absence/presence and rougher/smoother surface, respectively. **b**, ordination of averaged biofilms profiles analyzed as described in “a” with identical symbols. 2D stress was 0.11. Representative confocal micrographs of extreme biofilm phenotypes from the rougher surface are shown, each depicting an area of 295 µm^2^. **c**, vector overlay illustrating the correlations between biofilm parameters and the ordination axes, using the same ordination as shown in “’b”.

**Table 2.** Differences in biofilm profiles across experimental groups and time.

| Model | Source | Pseudo-F | P (perm) | Sq. root |
| --- | --- | --- | --- | --- |
| Treatment,<br>surface,<br>time | Tr. | 5.0 | 0.0033 | 7.6 |
|  | Su. | 13.9 | 0.0001 | 13.6 |
|  | Ti. | 3.6 | 0.0002 | 9.7 |
|  | Tr. x Su. | 1.5 | 0.1907 | 3.9 |
|  | Tr. x Ti. | 1.8 | 0.0427 | 7.6 |
|  | Su. x Ti. | 1.4 | 0.1728 | 5.0 |
|  | Tr. x Su. x Ti. | 1.3 | 0.1968 | 6.7 |
|  | Res. |  |  | 32.3 |

PERMANOVA was conducted to test the hypothesis of no differences in biofilm profiles with respect to 16 μM DHNA treatment (Tr), surfaces type (Su), and time (Ti). The table present pseudo-F ratio (Pseudo-F), permutation-derived *P* value [P (perm)] and square-rooted estimates of components of variation (Sq. root). Interactions between variables are indicated by x, while Res represents residual.

On the rougher surface, control and DHNA-treated biofilms were similar in the first week (though DHNA-induced aggregation was already visible), diverging in structure over time. Vector overlays highlighted correlations between biofilm parameters and ordination axes (**Fig. 4c**), with selected phenotypes illustrated by CLSM micrographs (**Fig. 4b**). DHNA-treated biofilms displayed larger aggregates and greater surface area coverage. While control biofilms evolved until day 3 before reverting to earlier profiles, DHNA-treated biofilms exhibited steady structural changes throughout the experiment.

### Simulation of biofilm models

The *in silico* model simulates biofilm dynamics on a titanium disk, tracking growth, nutrient concentration, and volume percentage over time (**Fig. 5a** – **5c**), as well as volume percentage relative to nutrient concentration (**Fig. 5d**). Simulation parameters are detailed in **Tab. 3**. Initially, biofilm was randomly distributed at the domain’s base, covering approximately 5 – 10% of the surface (**Fig. 5e**), while nutrients were uniformly distributed throughout (**Fig. 5f**). Unlike the *in vitro* experiments, the simulation continues until the entire domain is covered with biofilm and the nutrients are completely depleted.

**Figure 5.**
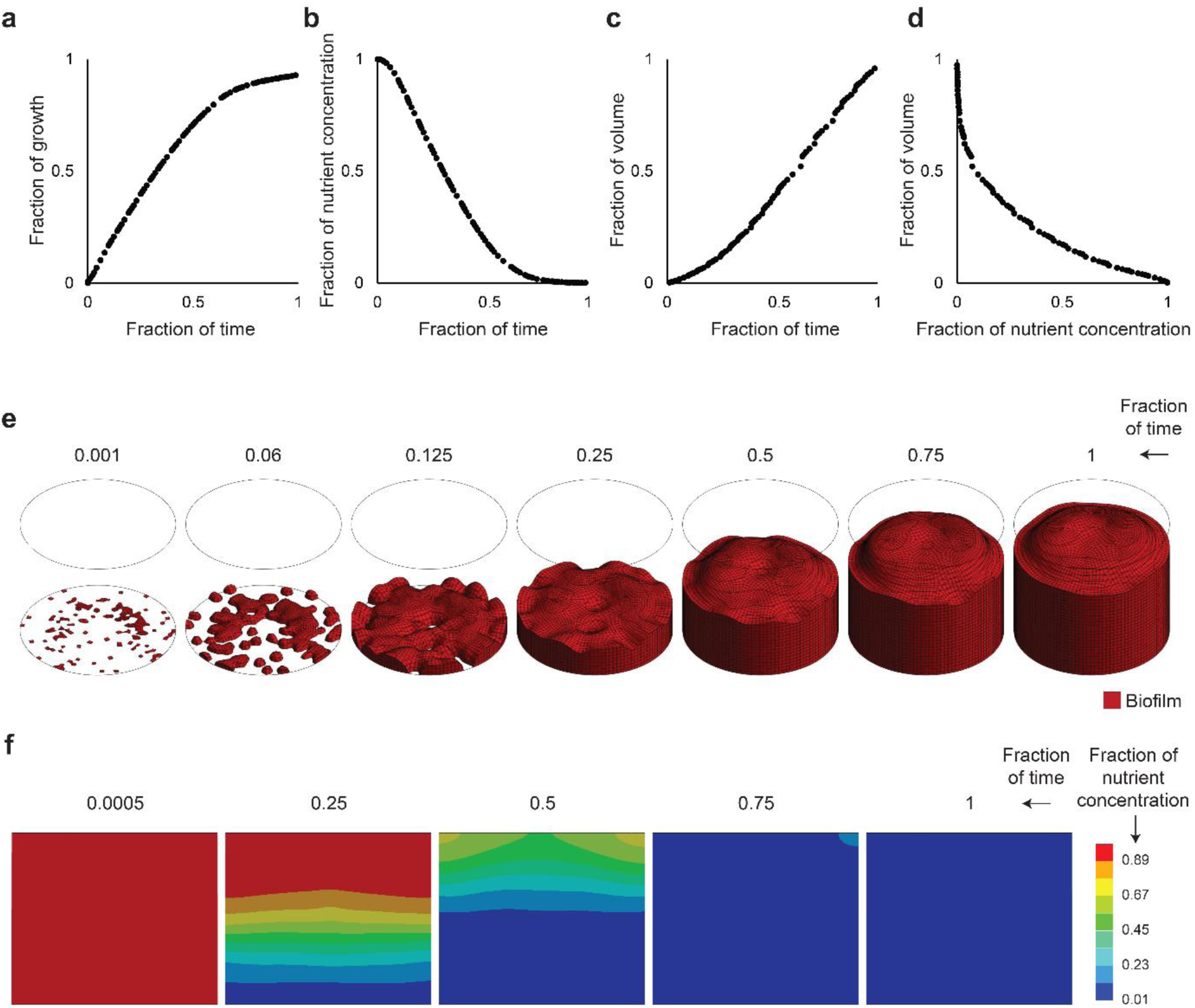
Simulation of biofilm development and nutrient utilization. **a**, growth over time. **b**, nutrient concentration over time. **c**, biofilm volume over time. **d**, biofilm volume relative to nutrient concentration. **e**, 3D snapshots of biofilm development over time. **f**, 2D snapshots of nutrient concentration over time. The cross section plotted is the middle section of the cylinder cut lengthwise. The parameters shown in this Figures are normalized. The nutrient concentration of one thus means a maximum of nutrients, not a specific amount, similarly is a simulation time of one not a fixed unit of time but rather the end of the simulated time.

**Table 3.** Simulation parameters for the biofilm model.

| Parameter | Value | Units |
| --- | --- | --- |
| Shear modulus $\mu$ | 3.35557 | Pa |
| Nutrient diffusivity $d$ | $8.6 \cdot 10^{-7}$ | $\text{m}^2 \text{day}^{-1}$ |
| Phase-field regularization parameter | $4.3 \cdot 10^{-8}$ | $\text{m}^2 \text{day}^{-1}$ |
| Growth factor | $8.6 \cdot 10^4$ | $\text{day}^{-1}$ |
| Half velocity constant | 2 | - |
| Consumption parameter | $8.6 \cdot 10^5$ | $\text{day}^{-1}$ |
| Growth parameter | $2.6 \cdot 10^6$ | $\text{day}^{-1}$ |

As the simulation progresses, the average value of the growth indicator variable approaches equilibrium, as shown in **Fig. 5a**. Nutrient depletion occurs alongside a simultaneous increase in growth (**Figs. 5a** and **5b**). Biofilm volume initially exhibits slow growth, transitioning to a linear increase towards the end of the simulation (**Fig. 5c**). When biofilm volume is plotted against nutrient levels, the relationship resembles *in vitro* growth patterns, particularly highlighting an initial dependence on ammonia (**Fig. 5d**). Over time, initially isolated bacterial colonies merge to form a continuous biofilm (**Fig. 5e**) while nutrients are entirely depleted (**Fig. 5f**).

## Discussion

Deciphering how specific metabolites influence the growth of periodontal pathogens within peri-implant biofilms is fundamental to resolve the persistent challenges posed by peri-implant infections. This study showed that the menaquinone precursor DHNA slightly stimulated *P. gingivalis* growth on implant-grade titanium, accelerating its transition to ammonia-independent growth, together with an increasing biofilm-covered area and aggregate size over time.

Exogenous naphthoquinones (NKs) and related metabolites, such as DHNA, have been shown to significantly impact bacterial growth and metabolism^22, 33, 65–67^, even at extremely low concentrations^68^. These compounds promote more efficient energy metabolism, enhancing bacterial survival and virulence, and also contribute to the development of pathologies. The effects of naphthoquinones (NKs) on biofilms are species- and condition-dependent. That, menaquinone supports *Staphylococcus aureus* growth and biofilm formation partly *via* SrrAB two-component regulatory system, is well known^69–71^. In contrast, menadione supplementation reduced matrix production and biofilm formation in menadione-auxotrophic *S. aureus* small-colony variants^72^, while phylloquinone significantly suppressed biofilm formation and virulence factor production in *Pseudomonas aeruginosa*^73^. In our preliminary work, we observed a complex network of NK-mediated interactions in the oral cavity involving *P. gingivalis*. Along with the effects of DHNA on *P. gingivalis* biofilms presented here, these findings suggest that electron transfer within oral biofilms, mediated by microbially excreted NKs, may play a crucial role in peri-implant diseases and could serve as a potential target for anti-biofilm therapies. Promising concepts for novel therapies include small-molecule inhibitors^46^, menaquinone analogs^74^, probiotic strains that compete with virulent species for the exogenous NK pool^28^, and bacteriophages targeting NK-releasing commensals^75–77^.

In parallel with monitoring *P. gingivalis* growth, we employed fluorometric estimation of ammonium ion concentration in *P. gingivalis* cultures and observed stronger ammonia depletion at an early growth phase, independent of treatment. This finding aligns with previous observations regarding the critical role of these ions in *P. gingivalis* growth on defined culture media^22^. The significant production of ammonia through amino acid catabolism and its role in environmental alkalization, which benefits *P. gingivalis*, is well known^78^. However, its role in anabolism and early population expansion remains largely understudied. The intense ammonia utilization observed during early growth in this study suggests that its specific role in early population expansion warrants further investigation.

Additionally, we analyzed the influence of implant surface roughness on DHNA-dependent biofilm growth. In previous studies, we adapted multivariate analyses from microbial ecology, such as MDS, PERMDISP, and PERMANOVA, to analyze biofilm profiles^60, 79^, and we also found them valuable for exploratory analysis here due to their simplicity and robustness^61, 62^. Here, we observed strong and, at times, unexpected surface-related biofilm effects. The increased colonization on rougher surfaces, which provides more attachment sites and sheltered niches, is a well-established phenomenon^80^, had been reported for *P. gingivalis*^48^, and was also confirmed in our study. Almost double the surface volume available for biofilm colonization was found for the total of the core surface (*Vvc*: 0.68 vs. 1.00 mL/m²) and the valleys (*Vvv*: 0.07 vs. 0.13 mL/m²) in the case of the rougher surface. However, on this surface, we observed an inverse volume dynamic in the DHNA-treated group, with a late increase, in contrast to the control group, which showed an early increase followed by a decrease and plateau. Additionally, we noted significant experimental variation, which may suggest the role of cell detachment in volume dynamics. Specifically, the DHNA group was characterized by extensive aggregation, which may promote detachment and, consequently, lead to a decrease in the observed volume. The molecular mechanisms underlying aggregation and how this is influenced by surface parameters require further investigation. A possible explanation could be potential defensive mechanisms against heme-related toxicity^81, 82^. Predicted detachment in dependence of surface roughness should be carefully studied, particularly in flow systems^83^ and through single-cell force spectroscopy^8^.

Our study also emphasized the importance of rigorous surface texture characterization in biofilm research. A more differentiated 3D surface texture parameter analysis has already been emphasized to improve the connection between biofilm development and surface properties^52^.

However, only the mean arithmetic height is commonly used in dentistry. In the present study, even though the two surfaces differ in the height of their features, the difference in the specific surface area available for biofilm adhesion seemed to be most important above all. We did not consider the large difference between the periodically recurring segments to be relevant for biofilm formation, as the size ratios are far above the expected bacterial size between ∼0.4 – 3 µm^3^ ^84^. The functional volume parameters offered additional insights into the available volume. Notably, the combined Vvc and Vvv values, representing the volume available for wetting the titanium surface (up to 0.75 µm³/µm² for the smoother surface and up to 1.13 µm³/µm² for the rougher surface), positively correlated with the observed biofilm volume. This relationship necessitates further investigation.

While our study provides valuable insights into the effects of DHNA on *P. gingivalis* biofilm growth, the study design is not free of limitations. The experiments were conducted in a rich medium, providing a standardized environment for analysis, although future studies can benefit from closely replicating the nutrient composition of saliva or serum *in vivo*^85, 86^. Additionally, the monospecies biofilm model offers valuable insights into *P. gingivalis* behavior, but a more comprehensive exploration of microbial behavior in the oral cavity can be obtained by incorporating polymicrobial models^87, 88^. Furthermore, the inclusion of a salivary pellicle and flow system in future studies would enable the simulation of initial adhesion and dynamic conditions^83^. Exploration of strain diversity, DHNA concentration, and physicochemical conditions, including varying oxygen levels, would enhance future understanding.

Incorporating all these aspects or dissecting their individual effects presents a significant experimental challenge. Therefore, in this work, we have initiated the development of *in silico* models to support experimentation and provide a more comprehensive understanding of *P. gingivalis* biofilm behavior. By integrating the empirical microscopic data on key biofilm parameters into our FEM-based simulation model, we have modeled dynamics of *P. gingivalis* biofilm growth. The successful alignment of the *in silico* model with *in vitro* growth and nutrient utilization highlights its potential as a cost-effective tool for studying biofilm development. Our *in silico* model can be further expanded to study the influence of surface textures on biofilm formation. Additionally, integrating additional species into model would enable the study of complex physical and metabolic interactions^68, 69^. By combining computational and experimental approaches, we can uncover how roughness-mediated microenvironments influence multispecies biofilm resilience. In the long term, this knowledge can guide the development of effective strategies for biofilm management and the optimization of surface designs in biomedical applications.

The key strength of our interdisciplinary approach lies in its ability to simultaneously capture the effects of growth factors and surface properties on *P. gingivalis* growth, while emphasizing the role of externally supplied naphthoquinones and ammonia in its early population expansion. In the future, our hybrid framework has the potential to increase experimental throughput and improve reproducibility of implant-associated biofilm research, as well as support the development of effective anti-biofilm strategies targeting metabolic interactions.

## Acknowledgement

Funded by the Deutsche Forschungsgemeinschaft (DFG, German Research Foundation) – SFB/TRR- 298-SIIRI – Project-ID 426335750 (KDN, MS, PJ, MSti & SPS). Additionally, PJ and MSti gratefully acknowledge the funding by the DFG, Research Unit 5250 (No. 449916462). We would additionally like to thank Andreas Winkel and Ines Yang for critical review of the manuscript, as well as Richard Werth, Diana Strauch and Sophie Ahlborn for excellent technical assistance.

