## Supplementary Information for "Biofilm development of *Porphyromonas gingivalis* on titanium surfaces in response to 1, 4-dihydroxy-2-naphthoic acid - a hybrid *in vitro* – *in silico* approach"

Short title: Effect of DHNA on *Porphyromonas* biofilm supplementary information

Rumjhum Mukherjee[1,2], Felix Klempt[3], Florian Fuchs[4], Katharina Doll-Nikutta[1,2], Meisam Soleimani[3], Peter Wriggers[3], Philipp Junker[3], Meike Stiesch[1,2], and Szymon P. Szafrański[1,2,$]

[1]Department of Prosthetic Dentistry and Biomedical Materials Science, Hannover Medical School, Hannover, Germany

[2]Lower Saxony Centre for Biomedical Engineering, Implant Research and Development (NIFE), Hannover, Germany

[3]Institute of Continuum Mechanics (IKM), Leibniz Universität Hannover, Hannover, Germany

[4]Department of Prosthodontics and Materials Science, Leipzig University, 04103, Leipzig, Germany

[$]correspondence to: Dr. Szymon P. Szafrański, Department of Prosthetic Dentistry and Biomedical Materials Science, Hannover Medical School Carl-Neuberg-Str.1 30625 Hannover, Germany;

**Table S1** Differences in biofilm profiles across experimental groups and time captured for each surface separately.

| Model | Surface | Source | Pseudo-F | P (perm) | Sq. root |
|---|---|---|---|---|---|
| Treatment, time | Smoother | Tr. | 5.0 | 0.0056 | 8.1 |
| | | Ti. | 1.9 | 0.0363 | 6.1 |
| | | Tr. x Ti. | 1.9 | 0.0405 | 8.6 |
| | | Res. | | | 24.6 |
| | Rougher | Tr. | 2.5 | 0.0544 | 8.0 |
| | | Ti. | 2.6 | 0.0034 | 13.1 |
| | | Tr. x Ti. | 1.3 | 0.1968 | 8.6 |
| | | Res. | | | 38.9 |

PERMANOVA was conducted for each surface to test the hypothesis of no differences in biofilm profiles with respect to 16 μM DHNA treatment (Tr), and time (Ti). The table present pseudo-F ratio (Pseudo-F), permutation-derived *P* value [P (perm)] and square-rooted estimates of components of variation (Sq. root). Interactions between variables are indicated by x, while Res represents residual.

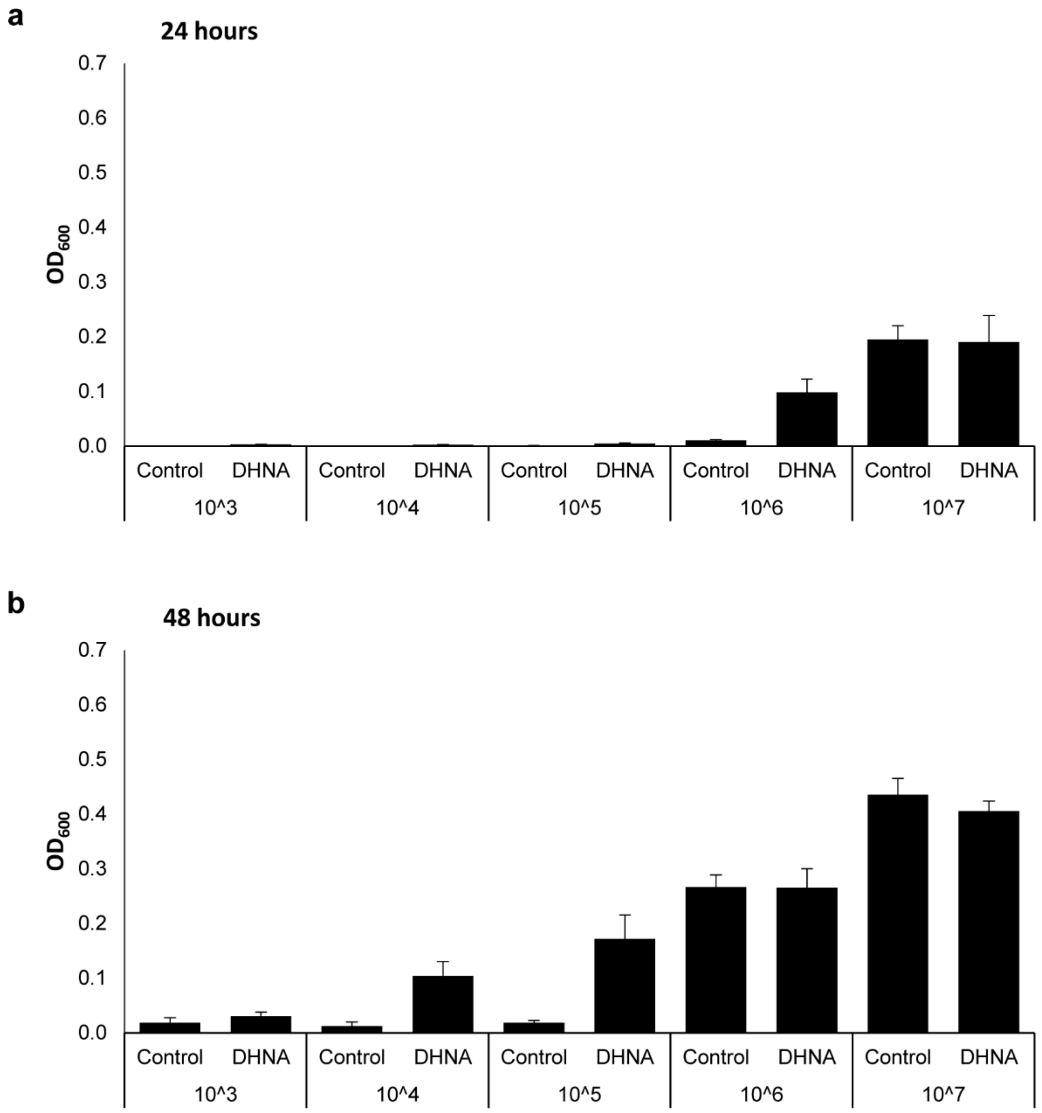


**Figure S1 DHNA enhances growth of *P. gingivalis* in a cell density-dependent manner over 24 and 48 hours.** Growth of *P. gingivalis* in the presence and absence of DHNA across a range of initial cell concentrations, measured by optical density at 600 nm ($OD_{600nm}$). Bacterial cultures were inoculated at serial dilutions ($10^3$ to $10^7$ cells/mL) and incubated for **a.** 24 hours or **b.** 48 hours. For each inoculum, $OD_{600nm}$ was recorded in cultures without and with 6 µM DHNA supplementation. Mean values represent at least five biological replicates, with SEM shown as error bars.

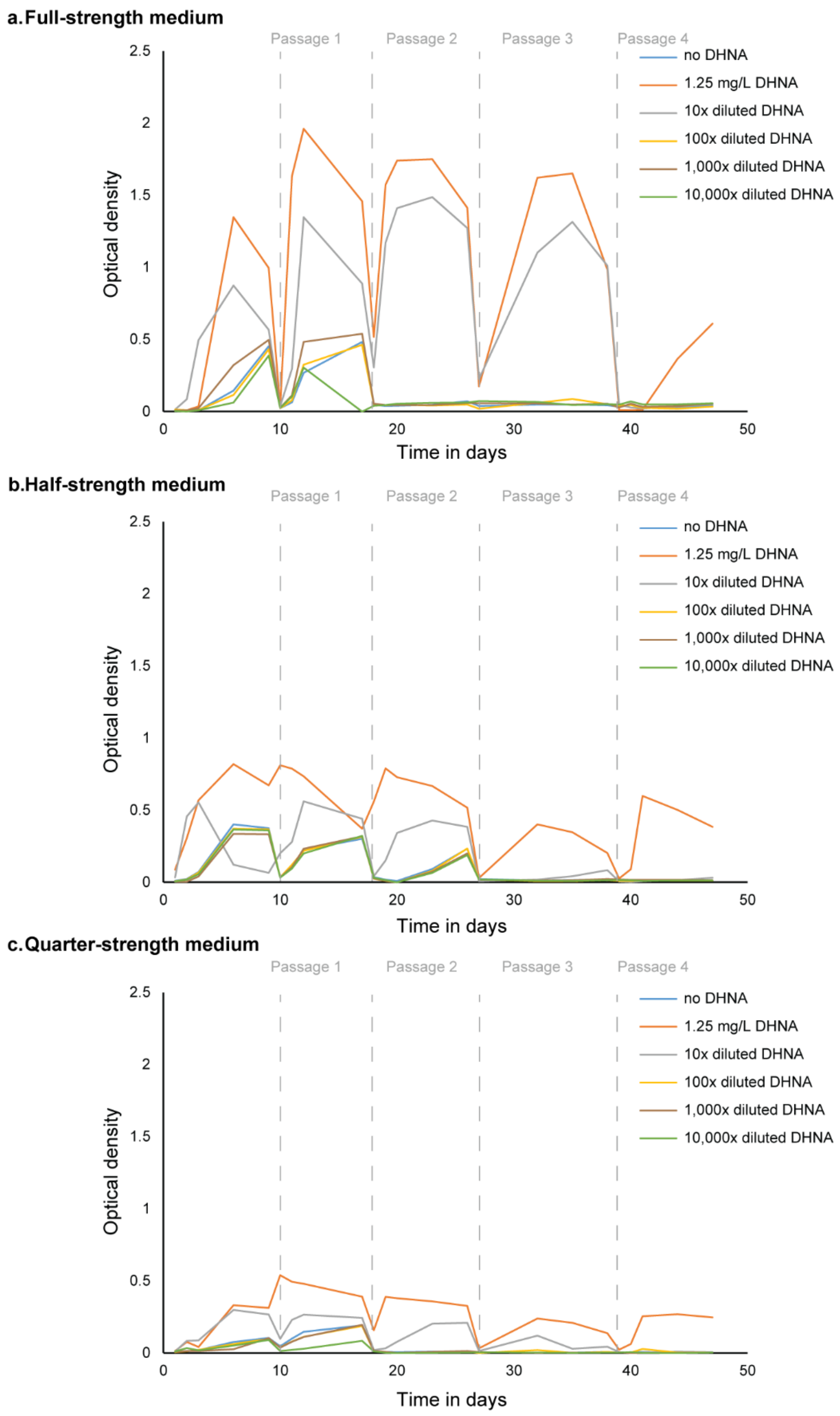


**Figure S2 Sustained *P. gingivalis* growth across nutrient conditions and serial passages supported by DHNA in a concentration-dependent manner.**
Growth dynamics of *P. gingivalis* cultured in **a.** full-strength, **b.** half-strength, and **c.** quarter-strength medium supplemented with varying concentrations of DHNA (1.25 mg/L to 10,000× diluted). Growth was measured as $OD_{600nm}$ and was monitored over 45 days, encompassing four serial passages. Each line represents a different DHNA concentration, including a control without DHNA. Mean values represent two biological replicates.
